# Insight into diatoms diversity at two European coastal sites (LTER-MC in the Mediterranean Sea and SOMLIT-Astan in the Western English Channel) using a DNA Metabarcoding approach

**DOI:** 10.1101/2022.07.01.498404

**Authors:** Mariarita Caracciolo, Cédric Berney, Benjamin Alric, Roberta Piredda, Adriana Zingone, Diana Sarno, Isabella Percopo, Sarah Romac, Florence Le Gall, Fabienne Rigaut-Jalabert, Anne-Claire Baudoux, Nathalie Simon, Nicolas Henry

## Abstract

Diatoms are among the most successful marine eukaryotic phytoplankton groups. Their diversity has been investigated in the world’s oceans through expeditions and observations carried out from the equator to the poles. Traditionally, diatom species have been distinguished based on morphological characters of their frustules, but high-throughput sequencing offers new, high-resolution data that can be used to re-examine spatial and/or temporal patterns of diversity. Here we investigated diatoms’ genetic diversity using metabarcoding (18S V4 rRNA gene) obtained along the years 2011 to 2013 at two coastal time series stations (SOMLIT-Astan and LTER-MareChiara) installed respectively off Roscoff in the Western English Channel and in the Gulf of Naples in the Mediterranean Sea. Diatom species pools detected were different, which fitted with previous observations and with our expectations, since these stations are installed in two contrasted pelagic habitats (permanently-mixed versus stratified in summer). However, this analysis also shows a pool of common ASVs among which some are persistent and dominant in both sites. The observed synchronous variations in relative read abundances of shared ASVs assigned to *Chaetoceros socialis, C. tenuissimu*s, *Cyclotella, Mediolabrus comicus* and *Leptocylindrus aporus* at the two geographically distant sites could indicate that internal controls of growth rate and sexual reproduction, rather that external environmental parameters are at work.

## Introduction

One of the most diverse and ecologically relevant groups of marine plankton is constituted by Bacillariophyta, better known as diatoms (Round et al., 1990; Werner, 1997; Kooistra et al., 2007; Behrenfeld et al., 2021). Diatoms are unicellular, almost exclusively free-living autotrophic organisms, with sizes ranging from approximately 2 µm to 1 mm (Mann et al., 2017). They are characterised by their silicified external cell wall, the frustule, whose size, thickness and ornamentation varies tremendously among species (Mann, 1999). Diatoms have colonised almost all illuminated aquatic environments. The vast majority of species are benthic, living attached to surfaces or gliding on sediment but they are also well represented in pelagic habitats (Round et al., 1990; Mann et al., 2017). In the ocean, they are by far the most successful eukaryotic phytoplankton group, not only in numbers of species but also in amount of biomass and primary production (Graham, Graham and Wilcox, 2009; Rousseaux and Gregg, 2013). They are responsible for roughly 40% of the marine primary production (Nelson et al., 1995; Field et al., 1998; Tréguer et al., 2018). Marine planktonic diatoms are abundant from the tropics to the poles, and most importantly in nutrient-rich coastal waters where they can form massive blooms (Arrigo et al., 2000; Assmy and Smetacek, 2009). The organic carbon that they synthesise at the surface in the euphotic zone fuels some of the most productive marine food webs and part of it is exported to depth as part of the biological pump (Tréguer et al., 2018). Diatoms thus participate in the provision of major ecosystem services on which humans depend socially and economically (Costanza et al., 1997; Hope et al., 2019).

Diatoms were among the first microscopic organisms to be detected in the sea, and, over the past two centuries, extensive occurrence data have been obtained through expeditions and observations carried in polar, temperate, tropical and equatorial waters (see Round et al. 1990, Estrada et al., 2016; Boltovskoy & Valentin, 2018; Sunagawa et al., 2020). Until recently, species occurrence and abundance were obtained using microscopy, which has traditionally been used for the description of diatom species whose frustules are highly differentiated. In this century, the use of genetic characters and of high-throughput sequencing (HTS) has shed light on the distribution and dynamics of species that are more difficult to identify with optical microscopy and has highlighted the widespread existence of cryptic species, which are not distinguishable based on morphological features (e.g. Sarno et al., 2007; Nanjappa et al., 2014). These techniques have also pointed to the widespread occurrence of small (nano-) diatoms (3-10µm) that can contribute significantly to spring blooms but also to carbon export (Leblanc et al., 2018; Arsenieff, Le Gall et al., 2020). Today, HTS enables extensive ecological studies through spatial gradients across the global ocean (i.e. Malviya et al., 2016; Piredda et al., 2018) and offers new, high-resolution data that can be used to re-examine temporal patterns of diversity and hypotheses about the factors and processes involved (Piredda et al., 2017; Caracciolo et al., 2022).

Here, we used a HTS method based on the metabarcoding approach to report on the genetic diversity of diatoms in 2 coastal sites with contrasting hydrological features but recurring annual sequences of plankton species (Zingone et al., 2019; Caracciolo et al., 2022): the SOMLIT-Astan and LTER-MareChiara time-series stations, located respectively in Roscoff (France, Western English Channel) and Naples (Italy, Mediterranean Sea, Gulf of Naples). We then investigated whether or not diatoms with identical 18S V4 rDNA sequences that thrive in both habitats have similar temporal dynamics. In particular, we were interested in discussing if species with identical sequences present at different sites follow the same seasonal cycle.

## Material and methods

### Sampling sites and sampling strategy

The SOMLIT-Astan and Long Term Ecological Research site MareChiara (LTER-MC; Fig. 1) are part of national and international monitoring networks: the SOMLIT (Service National d’Observation en Milieu Littoral) and PHYTOBS (Phytoplankton Observation Service) networks for SOMLIT-Astan and the Italian (LTER-Italy), European (ELTER) and International (ILTER) networks for LTER-MC. The general characteristics of these two sampling stations, which exhibit very different physico-chemical environmental and hydrological features, are described in detail by Guilloux et al. (2013) and Gac et al. (2021) for the SOMLIT-Astan station (also called SOMLIT-offshore), and by Zingone et al. (2019) for the LTER-MC station. Briefly, the SOMLIT-Astan station is characterised by permanently mixed waters (water column depth is 60 m) due to the intense tidal streams that characterise the French coasts of the Western English Channel (Pingree 1980). The heavy precipitation regime registered in winter in this region enhances river influx that consequently imports freshwater along with nutrient stocks in the marine environment (Tréguer et al., 2014). In contrast, the LTER-MC station, located in proximity of the 75 m isobath, at the transition between coastal mesotrophic water and oligotrophic blue water, is distinguished by a stratification period occurring from April to October (Ribera d’Alcalà et al., 2004), and very weak currents in particular during this period. Hence, while nutrients are never depleted in the permanently mixed waters of Roscoff, the establishment of a thermocline at Naples where incidental influx of nutrients occurs through runoffs from the watersheds can occasionally lead to limitations in nitrate and phosphate during summer (Ribera d’Alcalà et al., 2004).

**Figure 1.**
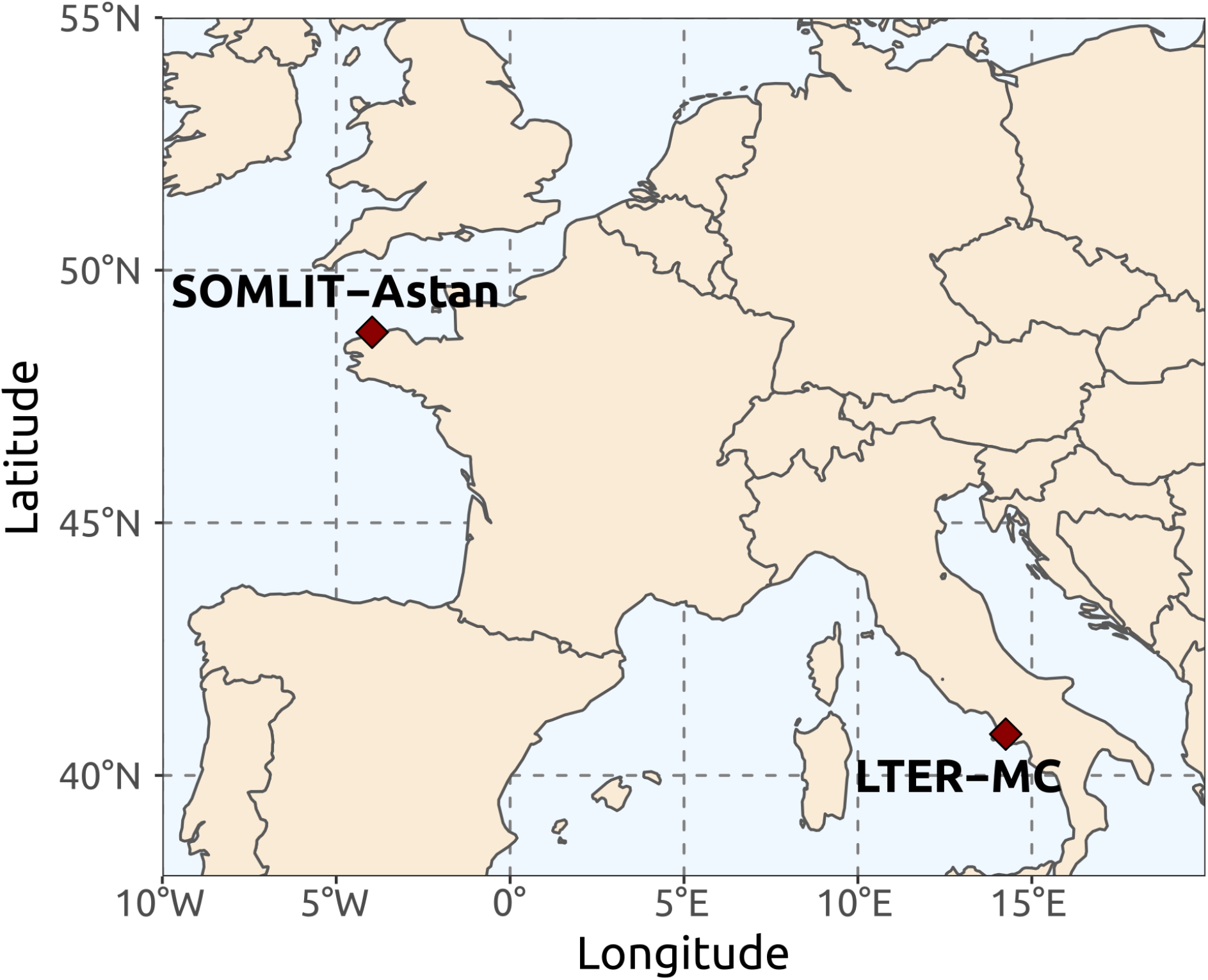
Sampling locations. The SOMLIT-Astan station (48°46’18’’ N – 3°58’6’’ W) is located 2.5 nautical miles off Roscoff in the Western English Channel and the LTER-MC station (40°49’N, 14°15’E) is located 2 nautical miles off Naples, in the Gulf of Naples, Mediterranean Sea.

At each station, seawater water samples were collected at the surface (1 m depth) from January 2011 to December 2013 using Niskin bottles (5 L). The temporal frequency of sampling was bi-monthly at the SOMLIT-Astan station (n = 69 samples collected systematically during neap tides), and more variable (0 to 4 times per month depending of the period) at the LTER-MC station (n = 40 samples). At each sampling event, a total seawater volume of 10 L and 20 L was collected at the SOMLIT-Astan station and at the LTER-MC station, respectively. Seawater samples were transported in the laboratory in Nalgene bottles and filtered onto 3 µm polycarbonate nuclepore membranes (47 mm, 5 L filtered) for SOMLIT-Astan and 1.2 µm cellulose ester membranes (47 mm, 3 L) for LTER-MC. Filters were then preserved in cryotube with 1.5 mL of lysis buffer (Sucrose 256 g.L^-1^, Trus 50 mM pH8, EDTA 40 mM) for SOMLIT-Astan and without lysis buffer for LTER-MC, and stored at −80°C until DNA extraction.

### Obtention of the metabarcoding data

The procedures used for DNA extraction and amplification of the 18S V4 region of the ribosomal operon are described respectively in Caracciolo et al. (2022) for the SOMLIT-Astan station and in Piredda et al. (2017) for the LTER-MC station. The eukaryote-specific primers used were TAReuk454FWD1 (5’-CCAGCASCYGCGGTAATTCC-3’, *Saccharomyces cerevisiae* position 565-584) and TAReukREV3 (5’-ACTTTCGTTCTTGATYRA-3’, *Saccharomyces cerevisiae* position 964-981) (Stoeck et al., 2010) for SOMLIT-Astan and TAReuk454FWD1 and V4 18S Next. Rev (5’-ACTTTCGTTCTTGATYRATGA-3’, *Saccharomyces cerevisiae* position 964-984) (Piredda et al., 2017) for LTER-MC. Sequencing was performed on Illumina MiSeq platforms (2×250bp) for both time series.Overall, 5,100,816 diatom reads were obtained for SOMLIT-Astan station and 12,524,142 for LTER-MC station. Raw sequences for SOMLIT-Astan are available at the European Nucleotide Archive (ENA) under the project id PRJEB48571.

The paired-end fastq files obtained from sequencing were demultiplexed. The sequence portions corresponding to the primers TAReuk454FWD1 and TAReukREV3 were trimmed from both SOMLIT-Astan and LTER-MC using Cutadapt v2.8 (Martin, 2011) and untrimmed reads were filtered out. This resulted in reverse reads from LTER-MC starting with three artificial nucleotides (TGA), but allows comparisons between the two sites. Then, forward and reverse reads were trimmed at position 220 and 210 respectively and reads with ambiguous nucleotides or with a maximum number of expected errors (maxEE) superior to 2 were filtered out using the function *filterAndTrim()* from the R-package *dada2* (Callahan et al., 2016). For each run, error rates were defined using the function *learnErrors*(), reads were dereplicated using the function *derepFastq()* function and denoised using the *dada()* function with default options before being merged. Remaining chimaeras were removed using the function *removeBimeraDenovo()*. Mixed orientated reads from the same sample were summed together and, in case of sequencing replicates, the readset with the highest number of reads was retained. Only amplicon sequence variants (ASVs) with at least two reads were retained. To assign taxonomy to ASVs, a native implementation of the naive Bayesian classifier method was used from the *assignTaxonomy()* function using PR2 v 4.13. For more details relative to the bioinformatic pipeline used to generate the ASV tables, see https://github.com/benalric/MetaB_Roscoff_Naples_diatoms. Finally, datasets corresponding to the SOMLIT-Astan samples (size fraction > 3 µm; years 2011, 2012 and 2013) and LTER-MC samples (size fraction > 1.2 µm; years 2011, 2012 and 2013) were pooled and ASVs observed in less than two samples were discarded. In order to take into account sequencing depth variability amongst samples, 24,974 reads were randomly sampled 10 times without replacement using the R function *rtk()* (rtk library). The rounded mean of the 10 values was reported as the number of reads in the rarefied ASV table. ASVs with 0 reads were considered as absent after rarefaction.

### Taxonomic reassignment of diatoms ASVs

ASVs assigned to Bacillariophyta with bootstrap confidence superior or equal to 99%, during the ASV table construction, were selected and reassigned, using a global pairwise alignment approach (usearch_global VSEARCH’s command), with an in-house diatom reference database. The reference database contains all diatom sequences from EukRibo (a manually curated rDNA 18S database, https://doi.org/10.5281/zenodo.6327891), as well as additional sequences from species or strains present at our sites but lacking in the database (for more details, see https://gitlab.sb-roscoff.fr/nhenry/rosko-naples-diatoms). ASVs inherited the taxonomy of the best hit or the last common ancestor in case of ties. ASVs with similarity scores below 90 % were discarded.

### Core diatoms

For each station, LTER-MC and SOMLIT-Astan, diatom ASVs detected in more than 30 % of the samples were defined as “core” (Magurran and Henderson, 2003; Krabberød et al., 2022). ASVs detected in at least one sample but not core for a given station were called “other”. The distribution of these 8 core-other-absent combinations were visualised with UpSet plots using the R library ComplexUpset (Lex et al., 2014; Krassowski, 2020).

### Phylogenetic analyses

Diatom ASVs defined as core in at least one of the two sites were aligned using MAFFT version 7.407 with the method L-INS-i (Katoh et al., 2013), together with the V4 region extracted from four Bolidophyceae sequences from EukRibo (AF167153, AF167156, AB275035 and KY980029) that were used as an outgroup. Aligned sequences were used to build a maximum likelihood phylogenetic tree using RaxML version 8.2.12 (Stamatakis et al., 2015) with 100 bootstrap replicates and using the GTR+CAT model of evolution.

## Results

### Different and shared diatom ASV pools in Roscoff and Naples

A total of 2,920 eukaryotic ASVs for 5,037,761 reads and 5,805 eukaryotic ASVs for 12,294,614 reads were retrieved for the SOMLIT-Astan station (Roscoff) and the LTER-MC station (Naples), respectively. Among these, 178 ASVs representing 924,969 reads and 235 ASVs representing 1,511,092 reads were assigned to diatoms for Roscoff and Naples, respectively. The Bacillariophyta represented 6.1 and 4.0 % of total ASVs richness in Roscoff and Naples, respectively. Altogether, our dataset contained 338 distinct diatom ASVs detected either in Roscoff or Naples and assigned to diatoms.

As expected in habitats as different as those represented by the SOMLIT-Astan station and LTER-MC station, a large proportion (78%) of the diatom ASVs detected along the years 2011 to 2013 were unique to either one or the other site (103 and 160 unique ASVs, i.e. 58 and 68 % of the ASVs at SOMLIT-Astan and LTER-MC respectively, Figure 2). They represent, however, only 42 and 25 % of the diatom reads. Most of these ASVs (76%) were identified as non-core (i.e. appearing in less than 30% of the samples examined in a given site).

**Figure 2.**
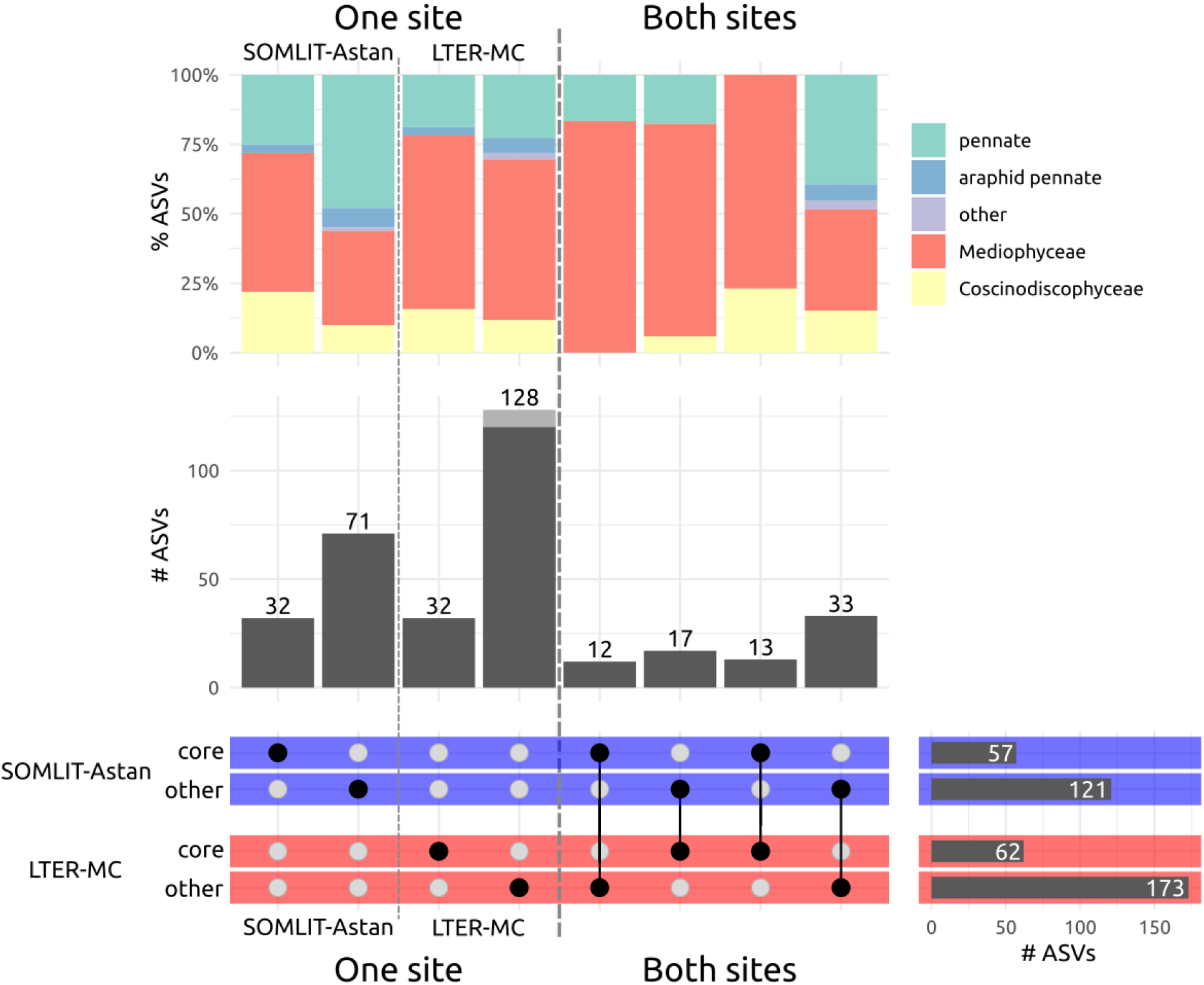
Comparative analysis (UpSet plot) of the pools of diatom ASVs detected at SOMLIT-Astan and LTER-MC between 2011 and 2013 (3 annual cycles). For each site, ASVs were clustered in 2 groups according to their persistence: ASVs detected in more than 30% of the samples were defined as “core” ASVs while the others were defined as “other”. The number of ASVs for each category is displayed in the bottom right panel. In the bottom left panel, the black dots indicate the intersections between the categories. A single black dot indicates ASVs unique to a category and two joined black dots indicate ASVs shared between two categories. For each intersection (unique and shared) the number of ASVs is shown in the barplots of the middle panel. The proportion of ASVs that is eliminated after rarefaction is indicated in grey. The taxonomic composition of each intersection is shown in the top panel.

A pool of 75 ASVs (42 and 32 % of the ASVs detected respectively at SOMLIT-Astan station and LTER-MC station) was shared between the two sites. Thirteen of these ASVs were identified as core at both sites and represent 27 and 32% of reads, at SOMLIT-Astan and LTER-MC respectively.

In both coastal sites, ASV pools were dominated by sequences assigned to the class Mediophyceae. In the pool of 13 ASVs that were shared and core in both stations, pennate diatoms were not detected. Conversely, pools of ASVs that were not identified as core in any of the 2 sites were enriched in pennate diatoms (both raphid and araphid) compared to all other pools of ASVs. In the pool of ASVs that were unique to the SOMLIT-Astan station, more than half of the ASVs were assigned to raphid pennate diatoms.

### Insight into the taxonomy and dynamics of core ASVs

We next explored the taxonomic assignments and seasonal dynamics of the 106 core ASVs detected in either the SOMLIT-Astan station or the LTER-MC station (Figure 3). Forty-two of these core ASVs (40%) were shared between the 2 sites.

**Figure 3:**
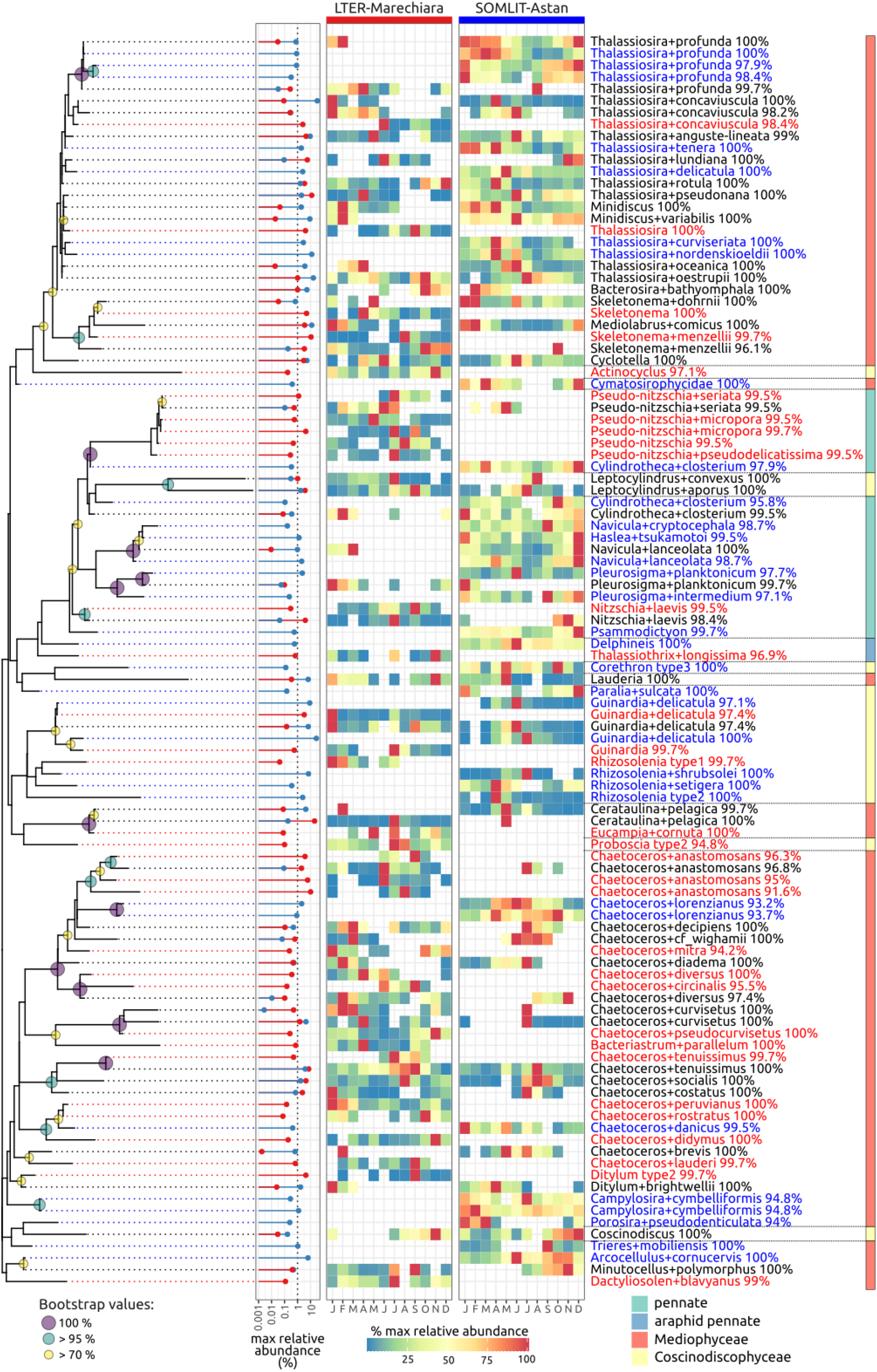
rDNA 18S V4 maximum likelihood phylogenetic tree of core diatoms detected during 3 years (2011-2013) at SOMLIT-Astan and LTER-MC stations. Bootstrap values above 70% are indicated by filled circles. Red (LTER-MC) and blue (SOMLIT-Astan) horizontal lines with a dot indicate the maximum relative number of reads as a proxy of abundance. The heat-map of the ratios between mean monthly relative abundances and maximum abundances indicates the seasonality. Taxonomic assignments and corresponding percentages of similarity are indicated in red for ASVs unique to the LTER-MC station, in blue for ASVs unique to SOMLIT-Astan station and in black for ASVs shared between the two sites. Note that because the tree topology was inferred only from the ca. 375 bp. long 18S V4 region, relationships between diatom lineages are not all recovered correctly (and only with low bootstrap support overall) and do not all match the current classification of diatoms.

At both coastal sites the Thalassiosirophycidae and Chaetocerotophycidae, two typically pelagic Mediophyceae sub-classes, appeared as the most diverse groups. These two sub-classes represented 51% of the ASVs detected as core in either one or the other station (28 ASVS for the Thalassiosirophycidae and 26 ASVs for the Chaetocerotophycidae). Within the Thalassiosirophycidae, the genus *Thalassiosira* was the most genetically diverse (17 and 13 ASVs detected respectively in the SOMLIT-Astan station and the LTER-MC station). The related genera *Minidiscus, Mediolabrus, Skeletonema, Bacterosira* and *Cyclotella* were also detected at both sites. Seventeen of these 28 ASVs were shared between the 2 sites and most of them were assigned to small-sized species (3-10 µm, *Thalassiosira profunda, Thalassiosira concavisuscula, Thalassiosira tenera, Thalassiosira pseudonana, Minidiscus* spp, *Mediolabrus comicus*). The sub-class Chaetocerotophycidae was mainly represented by genus *Chaetoceros* (with 11 ASVs shared between the two sites, and 3 and 12 ASV unique at the SOMLIT-Astan station and LTER-MC station, respectively). Members of the Cymatosirophycidae were also detected at both sites, with 6 ASVs (*Campylosira, Minutocellus*) at the SOMLIT-Astan station but only 1 ASV (*Minutocellus*) at the LTER-MC station.

The pool of core ASVs detected at either one or the other coastal sites also contained several members of the chain forming Rhizosolenioids and Leptocylindroids (formerly grouped in class “Coscinodiscophyceae”). The genus *Guinardia* was detected at both sites (with 1 shared ASV, and 2 additional unique ASVs at both sites). Core ASVs assigned to *Rhizosolenia* were only present at the SOMLIT-Astan station. Two ASVs assigned to the genus *Leptocylindrus*, and more precisely to *L. aporus* and *L. convexus*, were detected as cores in both coastal sites.

Raphid diatoms, a group that evolved adaptations to motility, were present in the pools of cores in both coastal sites. This group of diatoms was more diversified at the SOMLIT-Astan station, with 12 ASVs assigned to 9 genera (the truly pelagic *Pseudo-nitzschia* and the genera *Cylindrotheca, Delphineis, Haslea, Navicula, Nitzschia, Pleurosigma* and *Psammodictyon* which have benthic or tychopelagic affinities). At the LTER-MC station, 11 raphid pennate ASVs representing only 5 genera were identified among the cores (*Pseudo-nitzschia, Cylindrotheca, Navicula, Nitzschia* and *Pleurosigma)*. The genus *Pseudo-nitzschia* was more diversified at LTER-MC than at SOMLIT-Astan.

Focusing on the most persistent ASVs (presence in > 90 % of the samples) allowed us to detect large differences within the 2 sites. At the SOMLIT-Astan station, the ASVs assigned to the nanoplanktonic Thalassiosirales (*Cyclotella* sp., *Minidiscus variabilis, Thalassiosira concaviuscula, Mediolabrus comicus* and *Thalassiosira profunda*) and to the Cymatosirales (*Arcocellulus cornucervis*) were present in >90% of the samples analysed. The Thalassiosirales cited above were also detected at the LTER-MC station (respectively in 12, 67, 10, 47 and 7% of the samples studied) but *A. cornucervis* was not found. In the same way, ASVs assigned to *Cerataulina pelagica* and *Chaetoceros tenuissimus*, along with those assigned to *Leptocylindrus aporus* and *Chaetoceros socialis* were respectively present in all, or more than 90% of the samples of the LTER-MC station. These same ASVs were detected at the SOMLIT-Astan station but in lower proportions (in 1, 70, 36 and 57% of the samples, respectively).

Genetic diversity was found within some core species of both sites. For example, several ASVs were assigned to *Thalassiosira profunda* (2 ASVs identical to published sequences of strains identified as *T. profunda*) or *Guinardia delicatula* (4 different ASVs, one of which identical, and 3ASVs > 97% similar to published sequences of strains assigned to this species).

Most core ASVs showed pronounced seasonality at both sites (Figure 3, heat-maps, center panel) as expected in the 2 temperate environments studied. This was the case of ASVs assigned to members of the Thalassiosirophycidae and Chaetocerotophycidae which peaked in late winter, early spring or summer at both sites. Several ASVs assigned to *Thalassiosira profunda*, and an ASV assigned to *Mediolabrus comicus* were exceptions in this respect, since their maximum contributions were recorded in winter at both sites. In the SOMLIT-Astan station, some ASVs assigned to raphid pennates with benthic affinities (for example *Navicula, Pleurosigma, Psammodictyon*) had their maximum abundances in winter. A strong seasonal signal was also recorded at both sites for the large colony forming *Guinardia* and *Rhizosolenia*, or *Cerataulina*. Conversely, some core taxa had less pronounced seasonality in the SOMLIT-Astan station. This was the case for *Minidiscus variabilis* or *Campylosira cymbelliformis*.

### Seasonality and thermal niches of shared core and abundant ASVs in Naples and Roscoff

The seasonal variations exhibited by ASVs that were both core and abundant at both sites (presence in > 30 % of the samples and read counts above 10% of total eukaryotic read counts in at least one sample for both sites, Figure 3) were more carefully examined (Figure 4). For 5 of the 9 core and abundant ASVs (respectively assigned to *Chaetoceros socialis, C. tenuissimus, Cyclotella, Leptocylindrus aporus* and *Mediolabrus comicus)* patterns of variations observed along the 3 years studied were rather similar between the two sites. For example, for the ASV assigned to *Mediolabrus comicus* (100 % similarity to reference sequence), patterns of variations in relative read abundance were highly similar all along the 2011-2013 period, although the main peaks seamed to occur slightly earlier at SOMLIT-Astan (for example, in 2012, maximum contribution occurred in December in Roscoff and January in Naples). For the ASV assigned to *Cyclotella*, 2 seasonal peaks were observed each year, and at the same time in both stations. For all these species, given the differences in hydrology between the 2 habitats, environmental conditions that prevailed when their contributions were maximum differed greatly. As an example, for temperature, even if minima and maxima occur synchronously at LTER-MC and SOMLIT-Astan (minima in February/March and maxima in August, Figure S1), different ranges are observed along seasonal cycles (9.8 °C to 15.7°C off Roscoff and 13.9 °C to 26.6 °C in the Gulf of Naples). Hence, thermal niches observed for the same ASV with similar phenologies between the 2 sites differed by several degrees.

**Figure 4:**
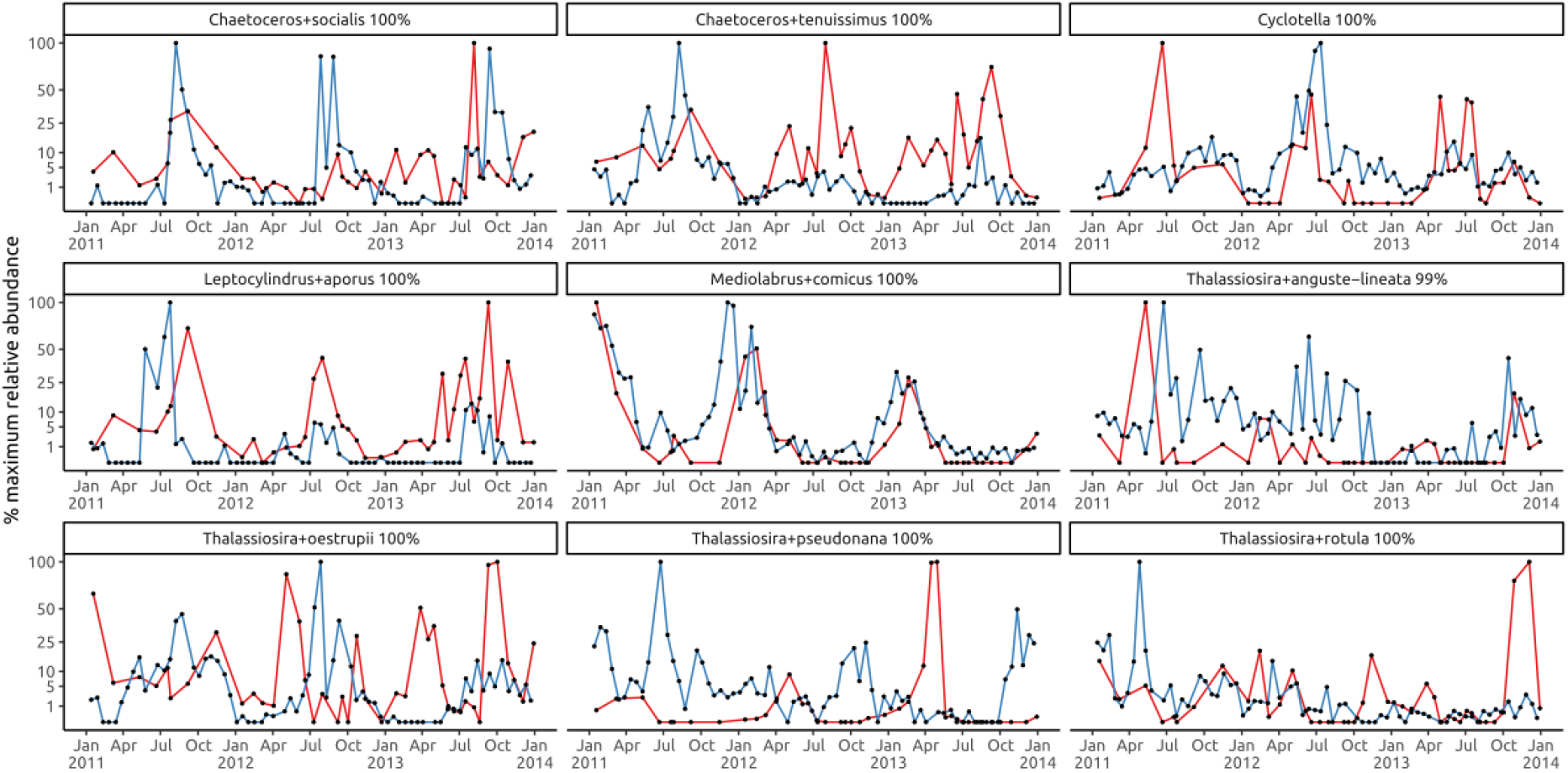
Abundance profiles of core and abundant diatom ASVs at LTER-MC and SOMLIT-Astan along the years 2011-2013. Abundance profiles are defined as the relative abundances (number of reads divided by the total number of eukaryotic reads) divided by the maximum relative abundance for each ASV at each site. Values, between 0 and 100%, are indicated in blue for SOMLIT-Astan and in red for LTER-MC.

The phenology of the 4 other dominating ASVs (respectively assigned to *Thalassiosira anguste-lineata, T. oestrupii, T. rotula and T. pseudonana*) differed between the 2 sites (Fig. 4).

## Discussion

The LTER-MC and SOMLIT-Astan time-series stations are installed in 2 different pelagic habitats (permanently-mixed *versus* stratified half the year) and biogeographical regions (Atlantic *versus* Mediterranean). Long-term records of phytoplankton dynamics indicate that diatoms dominate at both sites but the species pools and typical annual sequences of taxa differ substantially (Ribera d’Alcalà et al., 2004; Guilloux et al., 2013; Piredda et al., 2017; Zingone et al., 2019; Caracciolo et al. 2022). For example, the conspicuous late-spring/summer blooms in Naples are dominated by *Leptocylindrus aporus, Chaetoceros tenuissimus* and *Skeletonema pseudocostatum* (Zingone et al., 2019) while at Roscoff *Guinardia delicatula* is one of the greatest contributors to the blooms of that season (Sournia et al., 1995; Caracciolo et al., 2022). The relative high contribution of benthic and tychopelagic species to winter communities is also a distinctive feature of the SOMLIT-Astan station (Guilloux et al., 2013, Caracciolo et al., 2022), and more generally of tidally-mixed coastal habitats (Hernandez-Farinas et al., 2017). Previous direct comparisons of individual samples (1 sample from SOMLIT-Astan and 2 samples from LTER-MC from the water column, and 1 sediment samples from each site) produced using HTS (Nanjappa et al., 2014, Piredda et al., 2018) had also pointed to the marked individuality of these geographically distant sites. Our analysis of genetic datasets collected over 3 annual cycles which takes into account the highly seasonal character of plankton communities in temperate environments probably provides a more accurate image of the shared and unique lists of taxa/genetic types at the 2 sites.

The absence of ASVs in either one or the other site is always extremely difficult to interpret. It is also the case in our study based on datasets obtained from samples collected along several annual cycles. In our case, in addition, size fractions analysed differed between the 2 sites since plankton was collected onto 3 and 1.2 µm filters at SOMLIT-Astan and LETR-MC, respectively. The dataset from LTER-MC is thus probably enriched in sequences of pico- and nanoplanktonic species compared to that of SOMLIT-Astan. Yet, because cells smaller than the pore size, but also nucleic acids from broken cells of all sizes are always trapped on filters or stick to larger cells or particles, size fractionationation is never perfect (Chaffron et al., 2021; Caracciolo et al. 2022). The number of samples studied also differed between the 2 sites (69 and 40 samples examined in total for SOMLIT-Astan and LTER-MC, respectively). Hence, the fact that the total number of ASVs only found in LTER-MC (160 ASVs) was higher than in SOMLIT-Astan (103 ASVs) is difficult to interpret. One explanation would have been that the number of reads per sample is much higher for LTER-MC. After random subsampling at 24,974 reads, however, this difference is still observed with 152 and 103 ASVs for LTER-MC and SOMLIT-Astan respectively. Also, two different reverse primers were used for the PCR amplification steps for the 2 time-series stations (see material and methods) which probably led to differences in the overall relative abundances between the eukaryotic supergroups. In the future, analyses of datasets that are currently obtained on the long-term in a more concerted manner at time-series stations installed in different biogeographic regions and habitats will probably allow us to progress concerning questions regarding the ubiquity or restricted distribution of microscopic organisms. In fact, if Cermeno and Falkowski (2009, 2010) have elegantly shown that the “Everything is everywhere but the environment select’ famous statement fits with diatoms fossil records, doubts are still remaining. One of the reasons relates to the fact that cryptic species are common among diatom taxa, and in particular for the smallest ones.

In this study, one of the most important advantages of using datasets collected over several annual cycles was that it allowed us to determine pools of core ASVs (those present in > 30% of the samples examined) which can be considered as characteristic of the 2 respective habitats. A relatively high percentage of these ASVs (40%) that were core in either one or the other sites, were actually shared between the 2 sites. Pools of species and or genetic variants identified as core fitted with published lists of diatoms in the 2 sites, and included pelagic genera that are most common along the coasts of Europe (such as *Thalassiosira, Chaetoceros, Guinardia* and *Pseudo-nitzschia*, for which both shared and non shared ASVs were identified). For the genus *Guinardia*, which dominate spring blooms at SOMLIT-Astan, the sequence of the ASV that was unique for the SOMLIT-Astan site was 100% similar to sequences of strains isolated from that site (Arsenieff et al., 2019). Other core and abundant ASVs present in Naples and/or Roscoff and assigned to the genus *Guinardia* were more similar to a sequence of a *G. delicatula* strain isolated from the Gulf of Mexico (strain ECT3821, NCBI and Thériot et al., 2010). ASVs identical to that of *Leptocylindrus aporus* (which forms massive blooms in the Gulf of Naples) were also detected as core (and also abundant at both sites). Interestingly, an ASV 100% identical to *Leptocylindrus convexus* was also detected in both sites, while the species was not found in a previous examination of 2 samples collected at SOMLIT-Astan (Nanjappa et al., 2014). ASVs assigned to raphid pennate diatoms were detected at both sites. As expected, those associated to genera with benthic affinities were more diversified in Roscoff, and had higher contributions to total diatom reads. At the LTER-MC station, among ASVs assigned to pennate diatoms, only those assigned to the genera *Pseudo-nitzschia* and *Nitzschia* reached relatively high contributions to total diatom reads (Figure 3). One ASV assigned to *Nitzschia laevis* was actually identified as core in the 2 sites while the other one was unique to LTER-MC. The sequence of the latter ASV is highly similar (99.7%) to a sequence assigned to a dinotom (LC192341) described as *Durinksia* cf. *baltica* (Yamada et al., 2017). Dinotoms are dinoflagellates that harbour diatom endosymbionts. It is thus possible that the *Nitzschia* ASVs detected in both sites are actually endosymbiotic and it cannot be excluded that other ASVs, corresponding to other genera (such as *Cyclotella*, or *Chaetoceros*) known to be involved in these types of associations are endosymbionts also (Gavelis et al. 2018).

In our study, the most interesting result probably relates to the pool of 9 ASVs identified as both core and abundant at LTER-MC and SOMLIT-Astan. Five of these ASVs seemed to have highly similar phenologies at both sites, which appears as a surprising result at first glance because environmental conditions at the site studied differ considerably (see material and methods for a description of the 2 time-series stations) even if parameters such as day length, or temperature vary almost synchronously at the 2 sites. For *Chaetoceros socialis*, C. *tenuissimu*s, *Cyclotella, Mediolabrus comicus* and *Leptocylindrus aporus*, minima and maxima in reads abundance contribution are almost synchronous at both sites, which implies that their thermal niches differ greatly between the 2 sites. In fact, the observed dynamics in read relative abundance results from many different processes affecting either cell growth or cell losses *via* diverse mechanisms (for example, sedimentation, grazing, programmed cell death, internal clock, or infection by parasites). The synchronic variations in read contribution to total counts in such different and geographically distant sites could indicate that internal controls of growth rate and sexual reproduction, rather that external environmental parameters are at work (Cianelli et al., 2017; D’Alelio et al., 2010; Jewson et al., 2016; Longobardi et al. 2022). The 4 other species, identified as core and abundant in the 2 sites, may be more adapted to respond to natural environmental variability, and take advantage, for example, of pulses of nutrients.

To conclude, this study illustrates the importance of time-series stations and of the associated datasets produced in the long term for ecological studies. It also illustrates the importance of reference databases such as PR2 and EukRibo. Here, the final assignments of diatom ASVs used the manually curated EukRibo database. To the highly trustable subset of diatom reference sequences provided by EukRibo, we added a pool of reference sequences from strains observed at our time series stations and could thus accurately annotate most ASVs of our datasets. But of course, it will be important to go back to observations and experimentations in order to test the hypothesis that we proposed concerning, for example, the processes that control species phenologies in different environments.

## Acknowledgments

The authors would like to thank the captains and crew of the Neomysis research ship for their help during sampling at the SOMLIT-Astan station. They also thank A. Passarelli, F. Tramontano, M. Cannavacciuolo, and G. Zazo (SZN) and all the LTER-MC team for collaboration in sampling and data production, and the crew of the R/V Vettoria for assistance during the work at sea. We are grateful to the RCC for the maintenance of phytoplankton strains isolated from this station that served for the assignment of some of the genetic sequences. The Electron Microscopy facility of the SZN that helped with the analysis of diatom frustules is acknowledged. This study was supported by a PhD fellowship from Sorbonne University to MC, the French government research agency programs CALYPSO (ANR-15-CE01-0009), BIOMARKS (“Investissement d’avenir”, ANR-08-BDVA-0003) and OCEANOMICS (ANR-11-BTBR- 0008), the CNRS-INSU EC2CO CYCLOBS research project grant, the French Biodiversity Agency INDIGENE project (grant number OFB.20.0552), the Gordon and Betty Moore Foundation through the UniEuk grants GBMF5275 and GBMF8908 and the European Union program Assemble+ (Horizon 2020, grant agreement number 287589). We are grateful to the Roscoff Bioinformatics platform ABiMS (http://abims.sb-roscoff.fr), part of the Institut Français de Bioinformatique (ANR-11-INBS-0013) and BioGenouest network, for providing computing and storage resources.

## Data Availability Statement

The data that support the findings of this study are openly available in Zenodo at https://doi.org/10.5281/zenodo.6783055, Henry et al. 2022.

