## Supplementary material for "Insight into diatoms diversity at two European coastal sites (LTER-MC in the Mediterranean Sea and SOMLIT-Astan in the Western English Channel) using a DNA Metabarcoding approach": Figure S1

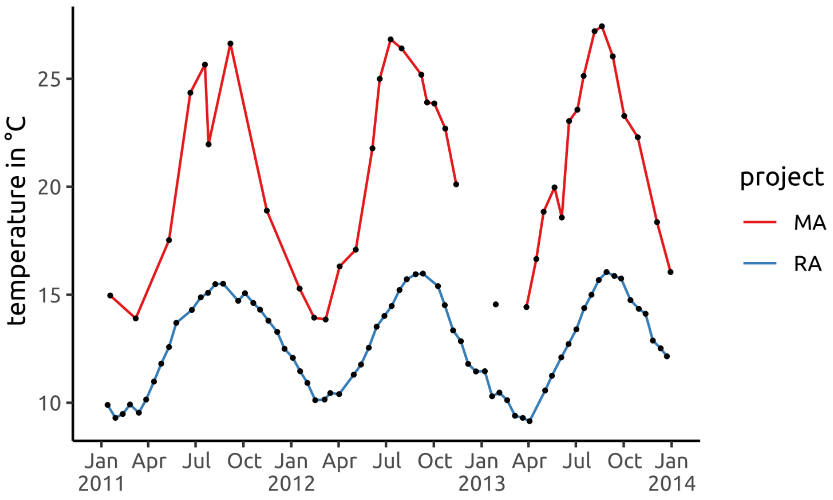


***Figure S1: Temperature profiles at LTER-MC and SOMLIT-Astan along the years 2011-2013.*** *Temperature values, in °C, are indicated in blue for SOMLIT-Astan and in red for LTER-MC.*
